# A microscopy reporter for cGAMP reveals rare cGAS activation following DNA damage, and a lack of correlation with micronuclear cGAS enrichment

**DOI:** 10.1101/2024.05.13.593978

**Authors:** Vivianne Lebrec, Negar Afshar, Lauren R. Davies, Tomoya Kujirai, Alexandra Kanellou, Federico Tidu, Christian Zierhut

**Affiliations:** The Institute of Cancer Research, Division of Cancer Biology, 237 Fulham Road, London SW3 6JB, UK; Laboratory of Chromatin Structure and Function, Institute for Quantitative Biosciences, The University of Tokyo, Bunkyo, Tokyo, Japan

**Keywords:** cGAS-STING, cGAMP, micronuclei, DNA damage

## Abstract

Cyclic GMP-AMP (cGAMP) synthase (cGAS) is the primary intracellular responder to pathogen DNA. Upon DNA-binding, cGAS generates cGAMP, which binds to STING, ultimately driving inflammatory signalling. Although normally silenced on self-DNA, cGAS can be activated during genotoxic stress. A universal by-product of these conditions are micronuclei, which accumulate cGAS, and which are therefore thought to be major cGAS activators. However, due to the inability to visualise cGAS activation in single cells, this hypothesis remains largely untested. Here we solve this question with an improved intracellular cGAMP reporter, which is compatible with microscopy, flow-cytometry and plate reader setups. Surprisingly, cGAS activation in response to multiple types of genotoxic stress is limited to a subfraction of cells and does not correlate with cGAS enrichment in micronuclei. Overall, our findings suggest a revised model of innate immune signalling in response to genotoxic stress, and introduce a novel and flexible tool with which to examine this model in future.

---

Most human cells express components of the innate immune system that allow rapid responses to infection^1^. A particularly important component of this response is cGAS, which senses pathogen DNA^2^. cGAS is activated upon DNA binding, generating the second messenger cGAMP^3,4^, which is a ligand for the ER-associated stimulator of interferon genes (STING)^2^. cGAMP engagement leads to the formation of STING oligomers, which translocate to the Golgi, where they activate NFkB and IRF3, leading to large gene expression changes and production of type I interferons such as interferon beta (IFNb)^2^. IFNb can then signal to the local environment through the STAT1/2 pathway, resulting in the induction of anti-pathogenic interferon-stimulated genes (ISGs)^2^. In addition, STING signalling can promote autophagy, cell death, senescence, proliferation and motility^5^. Moreover, cGAS and/or STING have also been suggested to have direct functions in genome stability^6^.

Although initially suggested to be exclusively cytoplasmic, cGAS is also present within the nucleus^7–9^ in most cells, raising the question why cGAS is not constantly activated by chromosomal DNA. We and others have reported that this is prevented by the inhibitory association of cGAS with nucleosomes, the building blocks of chromatin^5,8,10–16^. However, in response to the genotoxic stress from DNA damage and chromosomal instability, cGAS can be activated independently of infection^5^. The identity of the by-products of genotoxic stress that activate cGAS are currently unknown. However, a predominant model is cGAS that is activated in response to micronuclei. This is because micronuclei themselves are a universal consequence of genotoxic stress^17^. Formed by chromosome fragments, or mis-segregated chromosomes, micronuclei assemble fragile nuclear envelopes that are prone to rupture^17–19^, thereby allowing a dramatic enrichment of cGAS^7,20–23^. Whilst micronuclei are an attractive candidate for activating cGAS, micronuclei are composed of chromatinised DNA^24–26^, which should keep cGAS inactive. Additionally, RNA:DNA hybrids^27^ or mitochondrial DNA (mtDNA)^28–31^ were also indicated to promote cGAS activation.

The major obstacle to understanding how cGAS is activated following genotoxic stress is the lack of robust techniques for detecting cGAS activation in individual cells. Therefore, it is difficult to correlate cGAS enrichment on micronuclei with cGAS activation. Previous attempts to correlate micronuclei with cGAS employed mRNA sequencing of single cells isolated by microscopy-assisted laser microdissection^22^. These experiments indicated a limited induction of ISGs specifically in some micronucleated cells. However, ISG induction is a very indirect measurement of cGAS activation, as it depends on interferon, which is diffusible. Alternative approaches that have been used to identify cells with active cGAS include monitoring the nuclear translocation of IRF3^32^, and identifying cGAS molecules that are in close proximity with each other, which was proposed to reflect activation-associated cGAS dimerisation^33^. However, IRF3 nuclear translocation is an indirect measurement, based on an event that is transient, and can be initiated independently of cGAS^34^. Furthermore, we and others have shown that two cGAS molecules can be in close proximity when inactivated by nucleosomes^11,15^, implying that proximity does not necessarily indicate activation. Altogether, therefore, these approaches as limited in their ability to report on cGAS activation in a direct and reliable manner. As an alternative, a recent study employed Förster resonance energy transfer (FRET) to determine the presence of cGAMP within cells^35^. Whilst this approach provided a direct readout for the generation of cGAMP, the most upstream step in the pathway, several issues limited the use of this construct. For example, the FRET pair that seemed to be required was formed by a cyan fluorescent protein, mTFP, and an orange fluorescent protein, mKO2, which yields a fairly limited spectral overlap between donor emission and acceptor excitation. Furthermore, filters suitable for mKO2 are not part of standard microscopy setups. In addition, this reporter did not seem to function in the presence of endogenous STING. Here, we present a modified cGAMP FRET reporter system, which can function in the presence of endogenous STING, and is highly effective for microscopy-based analysis as well as in flow-cytometry and as a recombinant protein in solution. We used this reporter to investigate cGAS activation kinetics following genotoxic stress. Surprisingly, we found that, following genotoxic stress, cGAS is activated only weakly and in in only a subset of cells. In addition, we found no obvious contribution of the enrichment of cGAS on micronuclei to its activation.

## Results

### Design of an improved reporter for the detection of cGAMP

To investigate the cGAS activation state in single cells, we optimised a previously published genetically encoded FRET-based reporter, consisting of a fragment of STING sandwiched between a cyan and a yellow fluorescent protein^35^. In this approach, the conformation change that results from the engagement of cGAMP leads to an increased energy transfer from the cyan protein to the yellow protein (Fig. 1A). However, this reporter is not well-suited for microscopy, primarily due to the use of a sub-optimal FRET pair (mTFP1 and mKO2), which has a low spectral overlap, and an acceptor emission spectrum that is not readily compatible with standard microscopy filters. To improve this approach, we instead used an optimised FRET pair of very bright cyan and yellow proteins, mTurquoise2 (mTurq2)^36^ and YPet^37^, an approach that we previously used to generate a FRET reporter for DNA damage checkpoint function^38^. Importantly, these are also well-suited for standard microscopy filter sets. Using the cyclic dinucleotide binding domain (CDN) of mouse STING for the cGAMP-binding region as previously described^35^, we initially prepared two versions of this reporter, YPet-STING_CDN_-mTurq2 and mTurq2-STING_CDN_-YPet. We then electroporated plasmids encoding these into HeLa cells lacking cGAS^8^, either on their own, or together with cGAMP. cGAMP promoted a profound FRET increase for the mTurq2-STING_CDN_-YPet construct (Fig. 1B and C), but only a modest FRET increase for the YPet-STING_CDN_-mTurq2 construct (Fig. 1B). As expected, a mutated version of the reporter that does not stably bind to cGAMP^35^ did not show a FRET shift in the presence of cGAMP (Fig. 1D).

**Figure 1.**
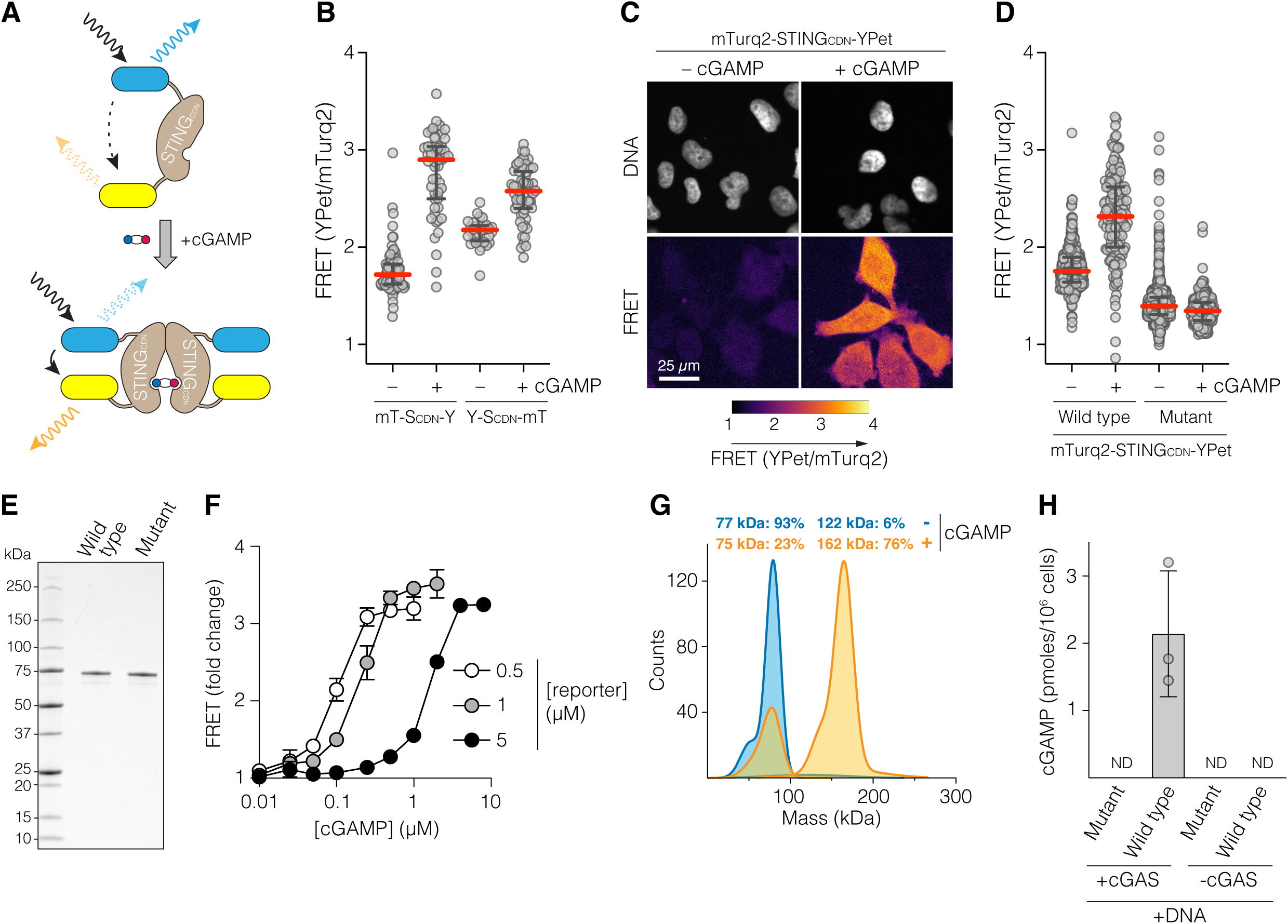
Design of an improved cGAMP reporter. (A) Schematic of FRET-based cGAMP detection. (B) Microscopy analysis of cGAS-disrupted HeLa cells electroporated with the indicated reporter constructs plus or minus cGAMP, and analysed after 48 hrs. Every dot represents a single cell. Red lines, medians; black lines, interquartile ranges (n= 34-83 for each condition). mT-S_CDN_-Y, mTurq2-STING_CDN_-YPet construct. Y-S_CDN_-mT, YPet-STING_CDN_-mTurq2 construct. (C and D) Microscopy analysis of cGAS-disrupted HeLa cells electroporated with the indicated reporter constructs plus or minus cGAMP, and analysed after 48 hrs. Representative images (C) and quantifications (D) are shown. Every dot represents a single cell. Red lines, medians; black lines, interquartile ranges (n= 157-817 for each condition). (E) Gel stained with Coomassie brilliant blue showing purified wild type and mutant reporters. (F) Plate reader based analysis of FRET generated by the indicated concentrations of reporter and cGAMP, normalised to unstimulated conditions. Averages and standard deviation from three independent experiments are shown. (G) Mass-photometry analysis of the oligomerisation state of the reporter with and without cGAMP. Monomer predicted molecular weight, 76 kDa. (H) Measurement of cGAMP production by cells +/- cGAS that were transfected with DNA. Cell lysates were analysed for cGAMP using the improved reporter, either in its wild type form, or in its cGAMP blind mutant form. Averages (bar) and standard deviation are shown. ND, not detected.

Endogenous STING is thought to form an inactive dimer in the absence of cGAMP, which undergoes conformational changes and oligomerisation upon cGAMP binding, with both steps being critical for downstream signalling^39–41^. To address the mechanism by which cGAMP engagement leads to an increase in FRET, and to characterise it in isolation, we purified the wild type and mutant reporters following overexpression in *E. coli* (Fig. 1E). The wild type reporter gave a concentration-dependent FRET shift over a wide range of cGAMP and sensor concentrations (Fig. 1F), whereas the mutated reporter did not (Fig. S1A). Mass-photometry analysis indicates that the apo-form of the reporter is primarily monomeric, whereas the cGAMP-engaged version is primarily dimeric, and does not form higher order oligomers (Fig. 1G). Similar results were also observed with native gel electrophoresis, suggesting that the engagement between the reporter and cGAMP is highly stable, and characterised by a low off-rate (Fig. S1B). Overall, we conclude that the cGAMP-engaged reporter is primarily dimeric, and that it is functional both when purified, and when present within cells.

### The new cGAMP reporter can be used in cell lysates, microscopy, and flow-cytometry

For bulk populations of cells, cGAMP production is often measured within cell lysates, either by mass-spectrometry, or by commercial ELISA kits. To determine whether the reporter can be used in a similar manner, we prepared lysates from HeLa cells with or without cGAS, and which had been transfected with DNA. Increased FRET was observed in lysates from cells expressing cGAS, but not in lysates from cells lacking cGAS (Fig. 1H). As expected, no response was observed with the mutant reporter (Fig. 1H). Overall, this indicates that the reporter can be used to determine the presence of cGAMP within cell lysates.

Although we were primarily interested in using this improved reporter for microscopy, we also tested whether it is suitable for flow-cytometry analysis. To do so, we transfected HEK293 cells, which lack endogenous cGAS, with plasmids driving expression of our improved cGAMP reporter. Subsequently, cells were either mock-treated, or incubated with 1 µM of the STING agonist diABZI^42^, harvested and analysed by flow-cytometry. FRET shift was observed within the treated population (Fig. S1C), indicating that our reporter can be used in flow-cytometry as well as in cell lysates.

### The new cGAMP reporter is highly sensitive for detecting cGAS activation in cells

We next generated wild type or mutant reporter-expressing versions of the non-cancerous breast epithelial MCF10A cell line, previously used to extensively characterise the cGAS response to DNA damage ^21,43^. Expression constructs were delivered on lentiviruses, and to prevent recombination during the lentiviral reverse transcription, we reduced the homology between mTurq2 and YPet by codon-optimising YPet for *E. coli*^44^ (Fig. S2A). Confocal live microscopy of cells that were either mock-treated, subjected to DNA transfection, or treated with either of two STING agonists, DMXAA^45–47^ or diABZI, revealed rapid responses to DNA and both agonists, specifically with the wild type reporter but not with the mutant reporter (Fig. 2A and B, Fig. S2B-D, Movie S1). For a cell line such as HeLa, which contains relatively high cGAS levels^8^, the intracellular concentrations are estimated to be in the range of ∼40 nM^48^, a concentration in which our cell line can discriminate between fine diABZI STING agonist levels (Fig. S2C). As expected, signal induced by DNA transfection was furthermore sensitive to chemical inhibition of cGAS^49^, verifying that FRET changes indeed report on cGAS activity (Fig. S2D). FRET signal was partially preserved after formaldehyde fixation, but to a lesser extent than when imaged under live conditions (Fig. S2E). Extensive washout did not dramatically change FRET for diABZI, although it did return to baseline levels for DMXAA, presumably reflecting differential off-rates of the two agonists (Fig. S2F).

**Figure 2.**
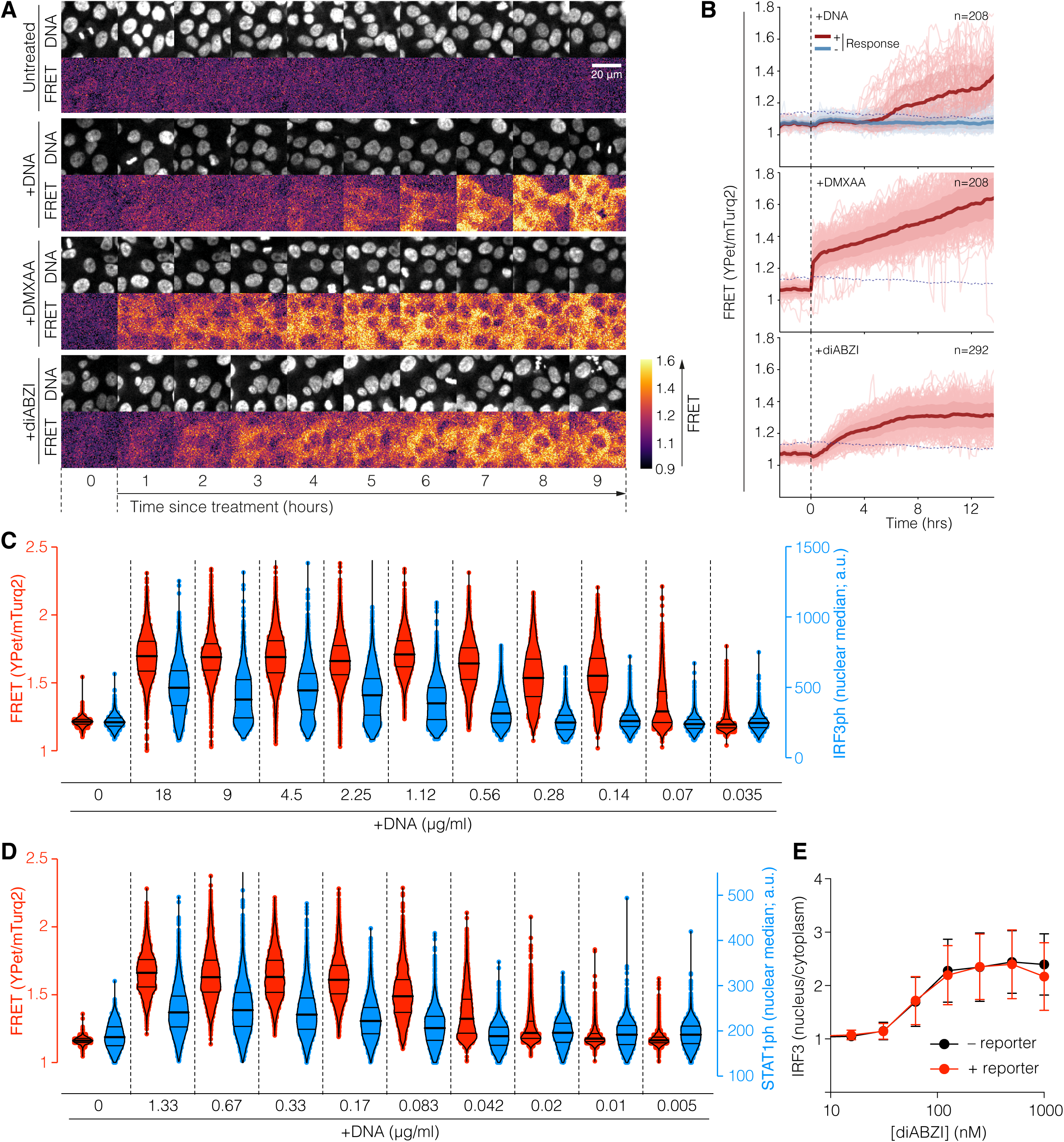
The new cGAMP reporter is highly sensitive for detecting cGAS activation in cells. (A and B) Time-lapse microscopy of MCF10A cells stably expressing the cGAMP reporter following mock-treatment or treatment with DNA transfection, DMXAA (10 µg/mL) or diABZI (1µM). Shown are images from representative views (A), and quantifications (B, dark lines, average; shaded area, standard deviation. Note that during this DNA transfection, not all cells received DNA, allowing us to divide cells into responders (transfected) and non-responders (untransfected) at the end of the imaging procedure. Dashed line marks the upper limit of FRET signal over time in control cells from the same experiment (see Fig S2B). (C and D) Comparison of cGAMP reporter FRET response (red, left y-axis) with nuclear signal of phospho-Ser396 of IRF3 (IRF3ph) (C), or phospho-Tyr701 of STAT1 (STAT1ph) (D) (blue, right y-axis). Cells were transfected with the indicated concentrations of DNA, and FRET was analysed after 24h (C) or 16h (D). Subsequently, cells were fixed and stained with the indicated antibodies. Individual datapoints are shown as dots and the general data are shown with violin plots (bold line, median; light lines, inter-quartile range). (E) Comparison of downstream signalling between MCF10A cells expressing the cGAMP reporter, and the parental cell line. Cells were incubated for 90 min with the indicated concentrations of diABZI, following which they were fixed and stained for total IRF3.

To compare the sensitivity of our reporter with established methods detecting downstream events, we treated cells with varying amounts of DNA transfection or diABZI, carried out live FRET imaging, and subsequently fixed cells and stained them with antibodies to either phospho-Ser396 of IRF3 (IRF3ph), or phospho-Tyr701 of STAT1 (STAT1ph). Cells from live FRET imaging were located in the follow up immunofluorescence (see Methods), allowing us to identify individual cells in both data sets. Overall, this indicated an increased sensitivity of our reporter when compared to these other methods (Fig. 2C and D). We also found that another potential live-cell reporter, GFP-tagged IRF3, could only be interpreted within a narrow window of time, due to the transient nature of IRF3 nuclear translocation - itself due to negative feedback control- and had overall reduced sensitivity compared to the improved cGAMP reporter generated in this study (Fig. S2G-I).

To test if the reporter might interfere with signalling downstream of STING, we treated the lentiviral cGAMP reporter cells and the parental cells lacking the reporter with a range of diABZI concentrations. Subsequently, cells were fixed and stained for IRF3, whose nuclear translocation was used as a proxy for cGAS signalling. No difference was observed between the two cell lines (Fig. 2E), indicating that the reporter concentration within the MCF10A cell line we have generated is unlikely to interfere with downstream signalling. Overall, we conclude that our new reporter is highly sensitive, and unlikely to interfere with downstream signalling when expressed at a suitable concentration. However, expression levels might require optimisation for each individual cell line.

### DNA damage and chromosomal instability lead to weak cGAS activation that manifests itself in only a minority of cells

With this optimised cGAMP reporter, we set out to investigate the cGAS response to DNA damage. We started with 15 Gy of ionising radiation (IR), similar to previously reported treatments to activate cGAS ^21^. Contrary to our expectations, only a minor fraction of cells showed evidence of cGAMP production, and the signal in each activating cell was generally lower than following agonist treatment and DNA transfection (Fig. 3A and B). FRET could be observed over a range of reporter expression levels, indicating that specific expression levels requirements for FRET after DNA damage did not exist (Fig S3A). Treatment with a cGAS inhibitor^49^ greatly reduced this response, and no signal increase was observed with the cGAMP blind reporter, confirming that this weak response to IR nonetheless depended on cGAS and cGAMP (Fig. 3C). Lower IR doses resulted in an even weaker response (Fig. 3D). Similar results were observed with doxorubicin, another inducer of DNA damage (Fig. S3B), and in the breast cancer-derived SUM149PT cell line (Fig. S3C-F). Time-lapse microscopy after irradiation furthermore revealed large temporal heterogeneity in the cGAS response, with some cells responding early, and some later, but again the majority of cells did not respond at all (Fig. 3E to G; Movie S2). DNA transfection at the end of the experiment confirmed that all cells had maintained competence for cGAS activation and reporter stimulation (Fig. S3G). Occasionally, we observed that cGAS activation in one cell was rapidly followed by appearance of FRET signal in neighbouring cells (Fig. 3H), suggesting cGAMP transfer via gap junctions, as was reported to occur following DNA transfection and viral infection^50^.

**Figure 3.**
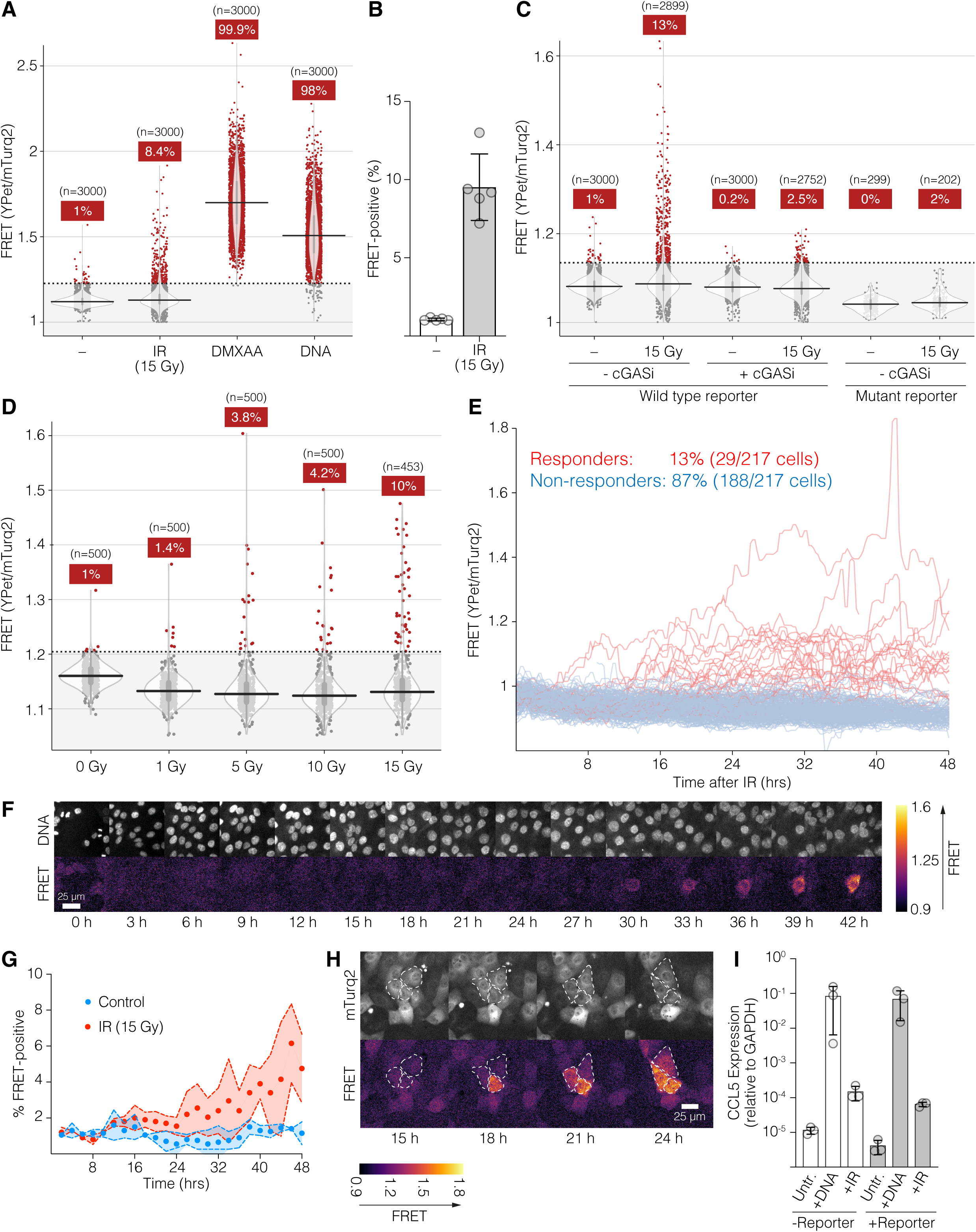
Genotoxic stress promotes only a weak cGAS response, which manifests in only a minority of cells. (A-H) Analysis of cGAMP reporter FRET response following the indicated treatments. For (A, C and D), dots represent values from individual cells. Violin plots show the general distribution, with the median indicated by a horizontal line, and the interquartile range indicated by a grey vertical line. Red dots indicate cells containing cGAMP (as defined by a FRET response higher than 99% of control cells). Numbers in red boxes indicate the percentage of cells containing cGAMP. (A and B) Comparison of cGAMP reporter FRET responses in untreated cells, cells treated with IR (15 Gy, analysis after 48 h), DMXAA (10 µg/mL, 2h treatment) or DNA transfection (1.33 µg/mL, analysis after 8h). Results from an individual experiment are shown in (A), and averages with standard deviation of five individual experiments for 15 Gy are shown in (B). (C) Analysis of the FRET response of cells expressing the indicated constructs, and subjected to the indicated treatments. Analysis was performed 48 h after irradiation. cGASi, treatment with 10 µM of the cGAS inhibitor G150. (D) Analysis of the FRET response of cells expressing the cGAMP reporter 48 h after irradiation with the indicated doses of IR. (E-H) Time-lapse analysis of the FRET response of the cGAMP reporter to 7Gy (F) or 15 Gy (E,G,H). (E) FRET traces of individual cells (n=217). (F) Images indicating cGAS activation in an individual cell. (G) Graph displaying the fraction of cells with FRET-response over time. Dots, averages; shaded area/dotted lines, standard deviation. (H) Example of cells activating the reporter in a small cluster. (I) Quantitative reverse-transcription PCR analysis of CCL5 induction by DNA (6h post transfection) and IR (48h after 15 Gy irradiation). Averages (bars) and standard deviation from three individual experiments are shown.

To confirm our single-cell analysis with a quantitative population-based assay, we analysed the induction of the ISG CCL5, a common target of the cGAS pathway. Whilst IR did indeed result in a modest increase in CCL5 expression, which was dependent on cGAS, it was nonetheless orders of magnitude less than the induction achieved by DNA transfection (Fig. 3I). Overall, we conclude that DNA damage provides only a weak stimulant for cGAS activation that manifests itself in only a minority of cells.

### Micronuclear accumulation of cGAS does not necessarily lead to its activation

Using γH2AX staining as a proxy for DNA damage, we established that cGAMP production was not generally correlated with the amount of persistent DNA damage that cells had received at a given dose of IR (Fig. S4A). However, rather than DNA damage itself, it is widely believed that enrichment of cGAS within micronuclei, derived from damaged chromosomes, drives cGAS activation^5,7,20–23^. To investigate this possibility, we performed live-cell FRET imaging after IR, followed by fixation and staining of cells with antibodies against cGAS. This analysis also allowed us to exclude the simple possibility that FRET signal was dependent on the intracellular cGAS concentration (Fig. S4B), or on its presence within/absence from nuclei (Fig. S4C). Within our datasets, we did indeed find examples of FRET-positive cells that contained cGAS-positive micronuclei (Fig. 4A, top, cell highlighted in yellow). Most frequently, however, we found that FRET positive cells did not contain cGAS-positive micronuclei (Fig. 4A, bottom, and Fig. 4B-C; Fig. S4D). In addition, we also found cells with cGAS-positive micronuclei that did not show evidence of cGAMP production (Fig. 4A, top, cell highlighted in green, and Fig. 4C).

**Figure 4.**
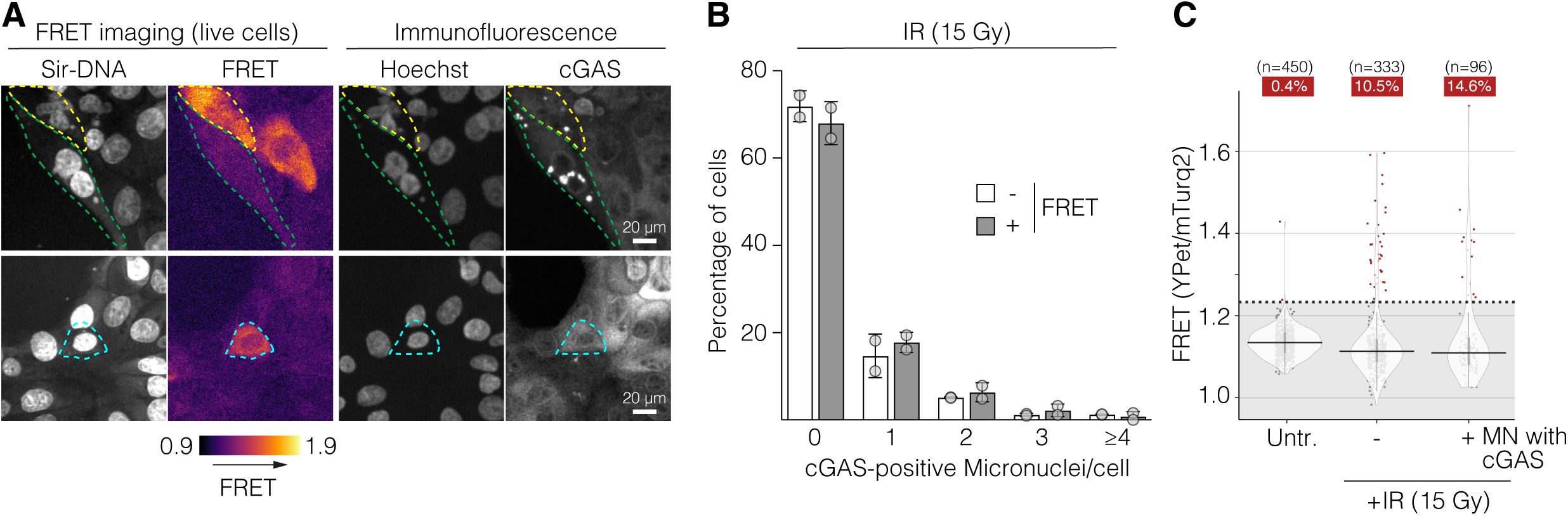
Activation of cGAS does not correlate with its micronuclear enrichment. MCF10A cells stably expressing the reporter were either mock-treated or irradiated with 15 Gy and imaged 48h later. (A-C) Analysis of correlative live-microscopy cGAMP reporter FRET imaging followed by immunofluorescence-based analysis of cGAS enrichment on micronuclei. (A) Representative examples of cells with micronuclear cGAS (upper panels) that either show reporter activation (cell outlined in yellow) or do not show reporter activation (cell outlined in green), as well as a cell with an active reporter that does not show evidence of micronuclear cGAS (lower panel, cell outlined in cyan). (B and C) Quantitative analysis of the correlation between cGAS-enriched micronuclei and cGAMP reporter FRET in cells treated with 15 Gy irradiation. (C) Cells exposed to 15 Gy radiation were categorised depending on the absence (–) or presence (+) of a cGAS+ micronucleus and according to the percentage of FRET+ cells compared to the non-irradiated control as in Fig 3A.

Since radiation promotes micronuclei formation by potentially complex processes, and because radiation may have other effects on cellular physiology, we also generated micronuclei by promoting chromosome mis-segregation independently of prior DNA damage. This was achieved by treatment with inhibitors of MPS1 (MPS1i), which leads to chromosome mis-segregation by weakening the spindle-assembly checkpoint^51,52^. We initially started by using 72 h of continuous treatment with 0.5 µM of reversine, conditions previously used to generate micronuclei^43,51^. However, we noticed that, in addition to micronuclei, this treatment caused extensive cell death and nuclear abnormalities (Fig. 5A). These were especially visible during live-cell imaging, but less so after fixation, presumably because fixation removed dying and dead cells (Fig. 5A and S5A-D). Since the nuclear abnormalities prevented us from investigating potential correlations between micronuclear enrichment of cGAS and its activation, we used pulses of various reversine concentrations, which resulted in the generation of cGAS-positive and cGAS-negative micronuclei, and at lower concentrations, this occurred in the absence of gross nuclear changes (Fig. 5A and B; Fig. S5B-D). At none of the concentrations used, even those that caused gross nuclear abnormalities, did reversine treatment result in frequent cGAS activation (Fig. 5C and D), indicating that, similar to IR, and doxorubicin, the genotoxic stress generated by MPS1 inhibition is not a very strong cGAS stimulant. Similar results were obtained with another MPS1i, AZ3146 (Fig. S5E). Correlative live FRET imaging followed by fixation and cGAS immunofluorescence revealed that, similar to IR, the large majority of cells that activated cGAS did not contain a cGAS-positive micronucleus (Fig. 5E). Furthermore, the majority of cells that contained a cGAS-positive micronucleus did not activate cGAS (Fig. 5E).

**Figure 5.**
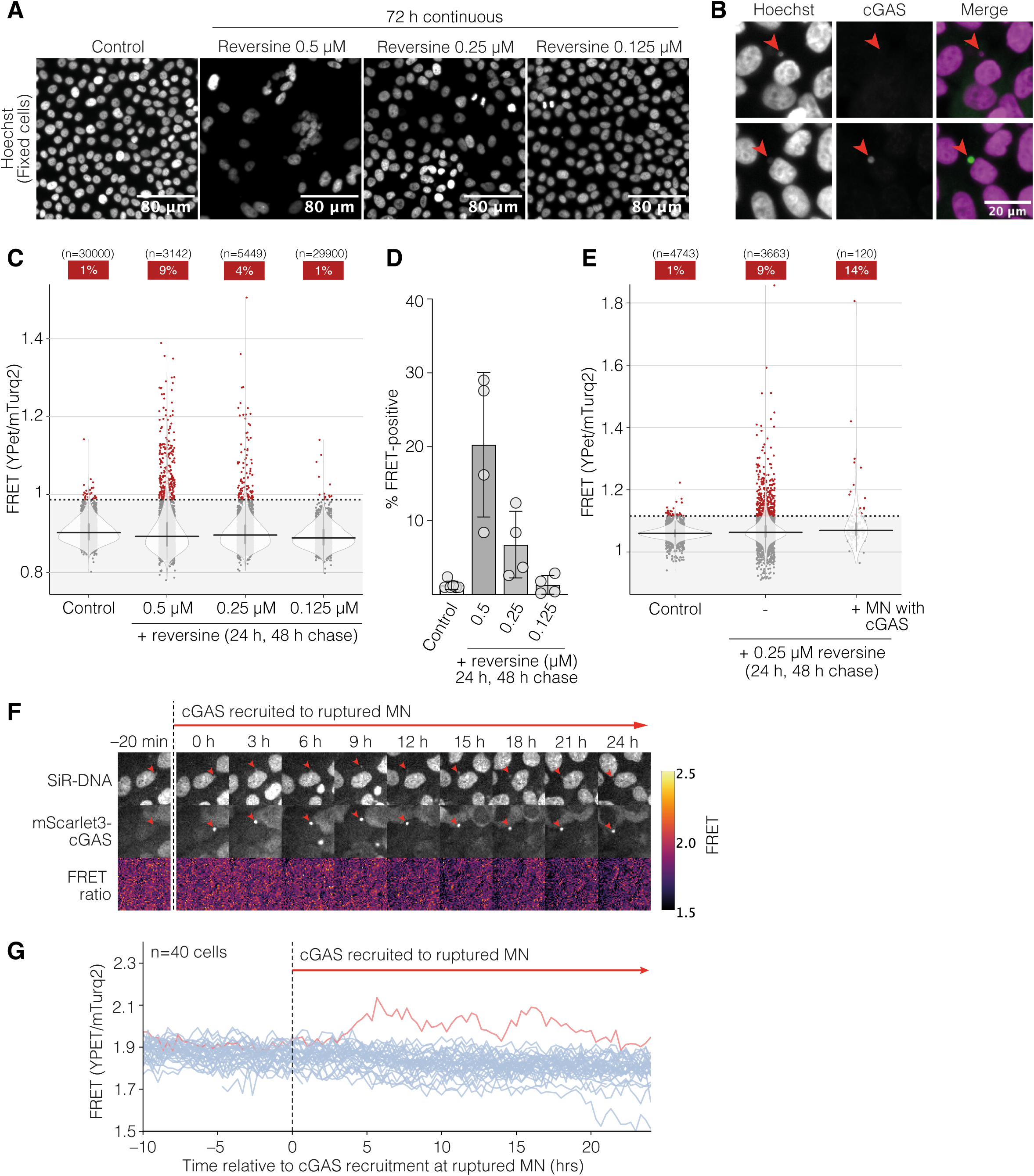
cGAS activation after chromosome missegregation does not correlate with micronuclear enrichment of cGAS. (A) Immunofluorescence analysis of MCF10A cells treated for 72h with the indicated doses of reversine (or DMSO control). Note the size and shape of nuclei treated with 0.5 µM. (B-E) MCF10A cells stably expressing the cGAMP reporter were treated with the indicated dose of reversine (or DMSO control) for 24h, following media change and FRET imaging or immunofluorescence 48h later. (B) Representative cGAS immunofluorescence stainings showing micronuclei with (lower panel) or without (upper panel) accumulation of cGAS in MCF10A cells treated with 0.25 µM reversine for 24 h, then released for 48 h before fixation. (C) Cells were plotted as in 3A. (D) Averages (bar) and standard deviation (error bars) of multiple experiments as in (C). (E) Analysis of correlative live-microscopy cGAMP reporter FRET imaging followed by immunofluorescence-based analysis of cGAS enrichment on micronuclei. Cells treated with a pulse of 0.25 µM reversine were categorised depending on the absence (–) or presence (+) of a cGAS+ micronucleus and depending on the percentage of FRET+ cells compared to the non-irradiated control as in Fig 4C. (F and G) Correlative time-lapse imaging of mScarlet3-cGAS enrichment on micronuclei and cGAMP reporter FRET treated with 0.5 µM reversine. (F) Images showing the behaviour of a representative cell. (G) Quantification. Cells were synchronised in silico to the time of cGAS recruitment to micronuclei. Each trace represents FRET variations over time for all quantified cells. Active cells are displayed in red, inactive cells in blue.

To directly assess whether micronuclear cGAS association correlates temporally with cGAMP production, we inserted a transgene expressing cGAS fused to the bright monomeric red fluorescent protein mScarlet3^53^ into the cGAMP reporter cell line (mScarlet3-cGAS). Time-lapse imaging after reversine treatment revealed that cGAS enrichment within micronuclei was not associated with subsequent cGAMP generation (Fig. 5F and G; Movie S3), even after prolonged periods of time. Overall, we conclude that during genotoxic stress, micronuclear enrichment of cGAS does not necessarily lead to its activation, and that cGAS activation can occur independently of micronuclei.

## Discussion

cGAS activation is a critical step during infection and genotoxic stress, regulating inflammatory responses to stimuli as diverse as bacterial and viral infection, irradiation, chemotherapy, genetic DNA repair deficiency and mechanical stress^2,5,54^. However, whilst it is highly likely that the DNA of pathogens is the major stimulant of cGAS during infection, it has been unclear how cGAS is activated during the sterile inflammation of genotoxic stress. Here we present an improved FRET-based reporter for cGAMP, which allowed us to address this question by monitoring the earliest step in the cGAS pathway, the DNA-dependent synthesis of cGAMP. This reporter can be used in microscopy as well as in flow-cytometry and plate reader-based assays. We showed that by careful generation of cell lines stably expressing this reporter, cGAS activation can be monitored in a manner that is more sensitive than previously described approaches. Because most other single-cell based approaches rely on events such as IRF3 phosphorylation or nuclear translocation, or expression of IRF3 target genes or ISGs - which can all occur independently of cGAS - our reporter is also more specific for cGAS activation than these other approaches. Moreover, the fact that FRET signal can be detected in flow-cytometry furthermore indicates that it may be useful to detect cGAMP production in a transgenic mouse model expressing the reporter. In the apo-state the reporter appears to be monomeric, and only dimerises in the presence of cGAMP, which is in contrast to STING constructs used for structural analysis^41,55^. At the moment it is unclear what the reason for this is, but we note that structural studies of STING are usually done under high concentrations (µM to mM and higher, in the case of crystallisation), which might promote dimerisation, whereas our analysis was done at very low concentrations (100 nM), which may be more representative of intracellular STING abundance. Alternatively, additional sequences close to the CDN of STING, which our construct lacks, might help dimerise apo-STING.

Careful cell line construction and experimental design will ensure robust results when using our cGAMP reporter. Cells with very low cGAMP levels might require low expression levels of the reporter. In addition, knockdown/knockout of STING may improve signal in cell lines with high STING expression levels, as endogenous STING could potentially compete for cGAMP. Low reporter expression levels also can ensure that the reporter does not reduce the pool of cGAMP that is available to stimulate endogenous STING. For example, in our experimental system, no obvious impact on signalling downstream of endogenous STING was observed (see Fig. 2E and Fig. 3I). It is unlikely that the reporter intercalates into oligomers of endogenous STING, as the reporter does not seem to intrinsically oligomerise (Fig. 1G and S1B), or shows evidence of aggregation inside cells (Fig. 1, 3 and 4). Potentially due to the lack of the membrane-domain, the reporter does not enrich at the Golgi, where cGAMP-bound STING oligomers are trafficked to^40,41,56–59^, and which ultimately leads to STING degradation in a manner regulated by the C-terminal domain^58,60–66^. This, combined with the high affinity of the STING CDN for cGAMP, and the potentially low cGAMP off-rate (Fig. S1B), suggests that the reporter may show delayed off-kinetics, which may however allow cGAMP production to be captured cumulatively, thereby facilitating detection of even weak or transient events.

In response to DNA damage and chromosomal instability, cGAS activation has so far primarily been investigated at the population level, suggesting a relatively uniform response of cells once they encounter genotoxic stress. However, using our reporter, we showed that cGAS activation during genotoxic stress induced by radiation, doxorubicin and chromosome mis-segregation is a rare event that moreover does not frequently result in very high cGAMP production. This was also supported by our own population-based analysis, which compared ISG induction between irradiated cells, and cells stimulated by DNA transfection (Fig. 3I). Revisiting previous work in this light suggests that low frequency cGAS activation may be a universal aspect of genotoxic stress^5,8,21,22,32,43,67^. Within the context of exposure to DNA damage agents, radiation and chromosome mis-segregation, our results suggest that rather than uniform cGAS-STING signalling throughout the exposed tissue, local centres of signalling may exist, which may expand by cGAMP transfer through gap junctions and cell migration (Fig. 6). This may be especially relevant within tumours, which frequently encounter genotoxic stress due to oncogene-mediated cellular dysfunction, chemotherapy and radiotherapy. Within this context, locally restricted signalling may limit immune responses, as well as impacting on other cancer cell behaviour^54^.

**Figure 6.**
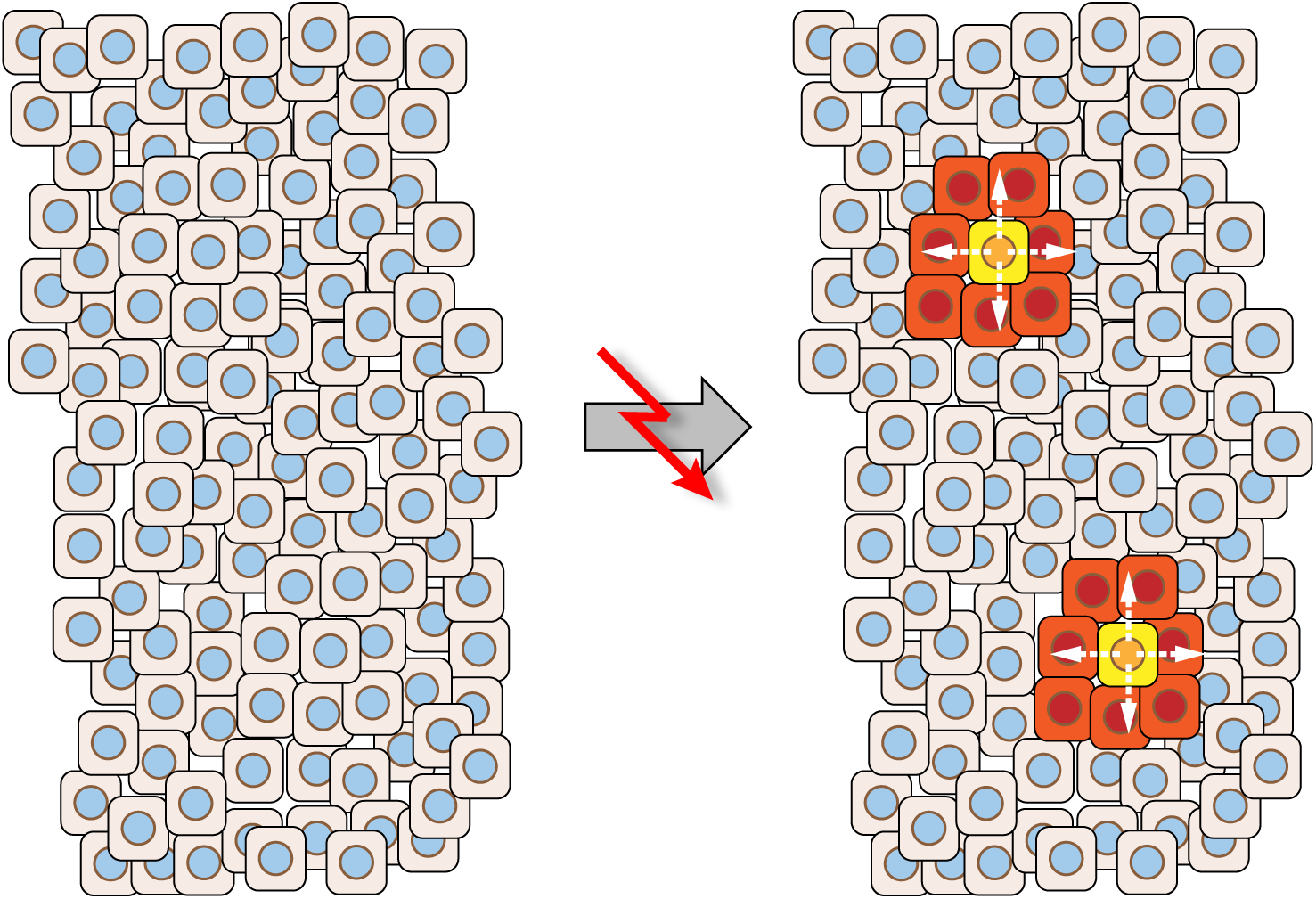
Model for cGAS activation within a tissue experiencing genotoxic stress. When a tissue experiences genotoxic stress, such as associated with radiotherapy and chemotherapy, our results suggest that cGAS-STING signalling and interferon production could be localised events, that could however, spread through cGAMP transfer and cell migration.

Testing a major hypothesis for how cGAS is activated by genotoxic stress, we have shown that the enrichment of cGAS on micronuclei, most commonly connected to cGAS activation, does not lead to cGAMP production. Similar results were recently reported using other means of detecting cGAS signalling^68^. While this may at first seem surprising, given the dramatic enrichment that cGAS can display on micronuclei, it is supported by previous evidence. Most importantly, micronuclei are composed of chromatinised chromosome fragments^24–26^, which we and others previously showed would keep cGAS largely inactive^5,8,10–16^. Even chromosome 21, the smallest human chromosome, composed of ∼45 million base pairs^69^, would contain a large excess of nucleosomes over the ∼50,000 cGAS molecules estimated to be present in a relatively high-expressing cell line such as HeLa^8,48^. Furthermore, previous work found only limited evidence for ISG induction in micronucleated cells^22^. Overall, while our work obviously does not exclude that micronuclear cGAS can be activated, it indicates that enrichment per se is not sufficient, and that other micronucleus-processing events may be required. In addition, our results show that cGAS can be activated in response to genotoxic stress independently of micronuclei. Whilst the nature of these structures is currently unclear, our improved cGAMP reporter can provide an excellent tool to identify these in future work.

**Figure S1.**
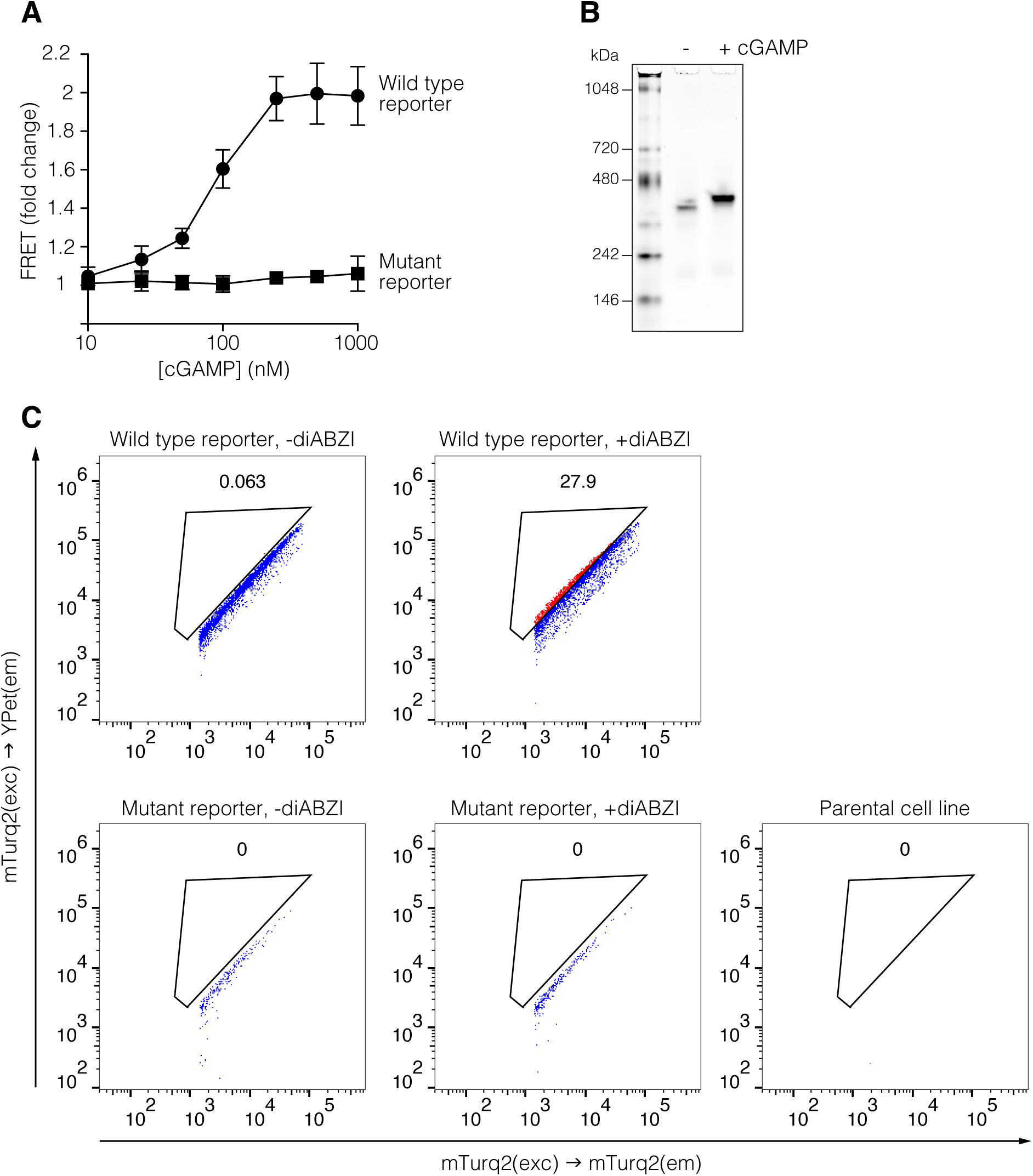
Further characterisation of the cGAMP reporter. (A) Plate reader based analysis of FRET generated by the indicated concentrations of wild type or mutant reporter and cGAMP, normalised to unstimulated conditions. Averages and standard deviation from three independent experiments are shown. (B) Native gel electrophoresis of the cGAMP reporter pre-incubated with or without cGAMP, stained with Coomassie brilliant blue. (C) Analysis of cGAMP reporter FRET in flow-cytometry. HEK293 cells were transfected with constructs promoting expression of the improved cGAMP reporter or its mutant cGAMP-blind version, and subsequently treated with 1 µM diABZI for 1.5 hrs. FRET positive cells are shown in red, FRET negative cells are shown in blue. Percentages of FRET positive cells indicated above gate.

**Figure S2.**
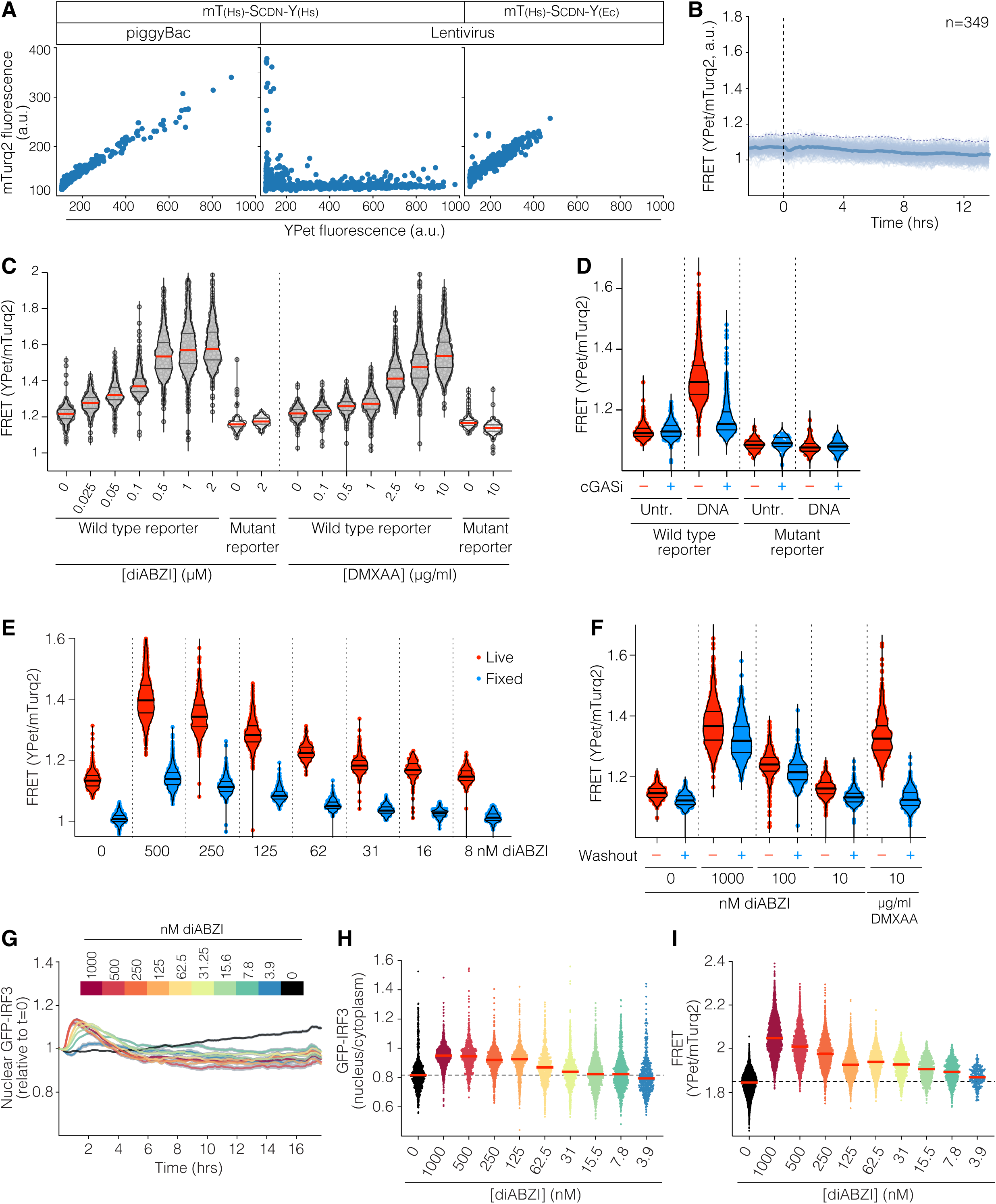
Further characterisation of stable cell lines expressing the cGAMP reporter. (A) Scatter plots of microscopy data from stable cell lines generated either by gene delivery based on piggyBac (left) or lentiviral particles, either before (middle) or after codon optimisation (right). Every dot represents the median signal from a single cell. (B) Quantification of cGAMP reporter FRET values over time for untreated cells. Light lines represent individual cells. Dark lines, average; shaded area, standard deviation. The dotted line going left to right represents 99% of the data. The dotted vertical line indicates the point at which drugs or DNA were added for the graphs shown in Fig. 2A and B. (C) FRET response of cells expressing either wild type or mutant reporter following 15h incubation with the indicated concentrations of diABZI or DMXAA. Dots represent individual cells (n=300-1400 cells for each condition). Red horizontal lines, median; black horizontal lines, interquartile range. (D) FRET response of cells expressing either wild type or mutant reporter 8h after mock-transfection or transfection with 1.33 µg/ml DNA, with or without 10 µM cGAS inhibitor (cGASi) G150. Dots represent individual cells (n=43-600 cells for each condition). Bold horizontal lines, median; light horizontal lines, interquartile range. (E) Comparison of FRET signal of cells imaged under live conditions (red) or following fixation (blue). Cells were incubated for 15h with the indicated concentrations of diABZI, imaged, fixed and imaged again. Dots represent individual cells (n=127-1600 cells for each condition). Bold horizontal lines, median; light horizontal lines, interquartile range. (F) Comparison of FRET signal of cells incubated for 15h with the indicated concentrations of diABZI or DMXAA, imaged either before (red) or after 60 min of washout (blue). Dots represent individual cells (n=200-1350 cells for each condition). Bold horizontal lines, median; light horizontal lines, interquartile range. (G-H) MCF10A stably expressing GFP-IRF3 were treated with the indicated dose of diABZI, and imaged every 5 minutes for 18h. (G) Mean ± SEM of nuclear IRF3 normalised to t0. (H) Column scatter plot of GFP-IRF3 nucleus/cytoplasm ratio at 60 min for every cell from (G). Each dot is a measurement from a cell, red horizontal bar is the population median. (I) In parallel to G-H, MCF10A cells stably expressing the cGAMP reporter were treated with the same doses of diABZI STING agonist. FRET in each cell was measured by imaging 18 h after stimulation with diABZI.

**Figure S3.**
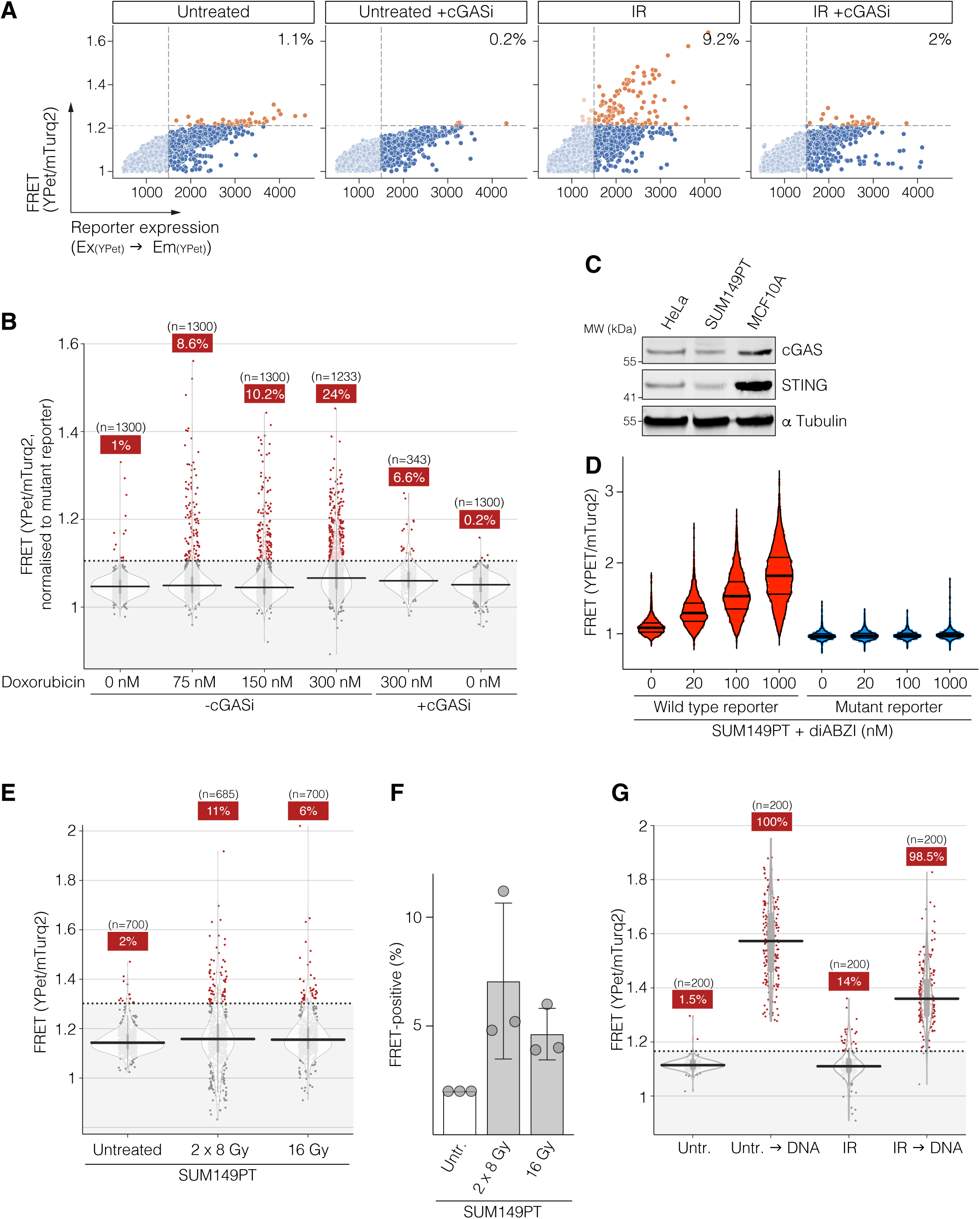
Further characterisation of the cGAS response to DNA damage. (A) Scatter plots indicating the relationship between cGAMP reporter expression levels (x axis) and FRET (y axis). Cells were either untreated, or treated with 15 Gy of IR in the presence or absence of 10 µM cGAS inhibitor G150. After 48 h, cells were analysed by live-microscopy. Cells of a YPet intensity below 1500 were considered too low-expressing for reliable cGAMP detection. (See Method section “Image processing and data analysis”.) (B) cGAMP reporter FRET response following doxorubicin. Cells were incubated with the indicated concentrations of doxorubicin, either on its own, or in the presence of 10 µM cGAS inhibitor G150. FRET imaging was performed after 48 h. Due to fluorescence from doxorubicin affecting the baseline of measured FRET, measured FRET values from each cellswere normalised to the average FRET value of MCF10A stably expressing the mutant cGAMP-blind reporter that were treated with the same dose of doxorubicin and imaged in parallel. (C) Western blot comparing cGAS and STING expression levels between Hela, SUM149PT and MCF10A cell lines. (D) FRET response of SUM149PT cells expressing either wild type or mutant reporter following 16 h incubation with the indicated concentrations of diABZI. (E-F) SUM149PT cells were either untreated, treated with 16 Gy of IR on day 1, or treated with 8 Gy of IR on days 1 and 2. Cells were imaged on Day 3. (F) shows the percentages of cells with detected FRET from three independent experiments. (G) MCF10A cells stably expressing the cGAMP reporter were either untreated, or treated with 15 Gy of IR. After 40h, each of the samples was either transfected with 1.33 µg/ml DNA or left untransfected. Cells were imaged and for FRET ratio measurement 8 h later. For (B), (E) and (G), dots represent values from individual cells. Violin plots show the general distribution, with the median indicated by a horizontal line, and the interquartile range indicated by a grey vertical line. Red dots indicate cells containing cGAMP (as defined by a FRET response higher than 99% of control cells). Numbers in red boxes indicate the percentage of cells containing cGAMP.

**Figure S4.**
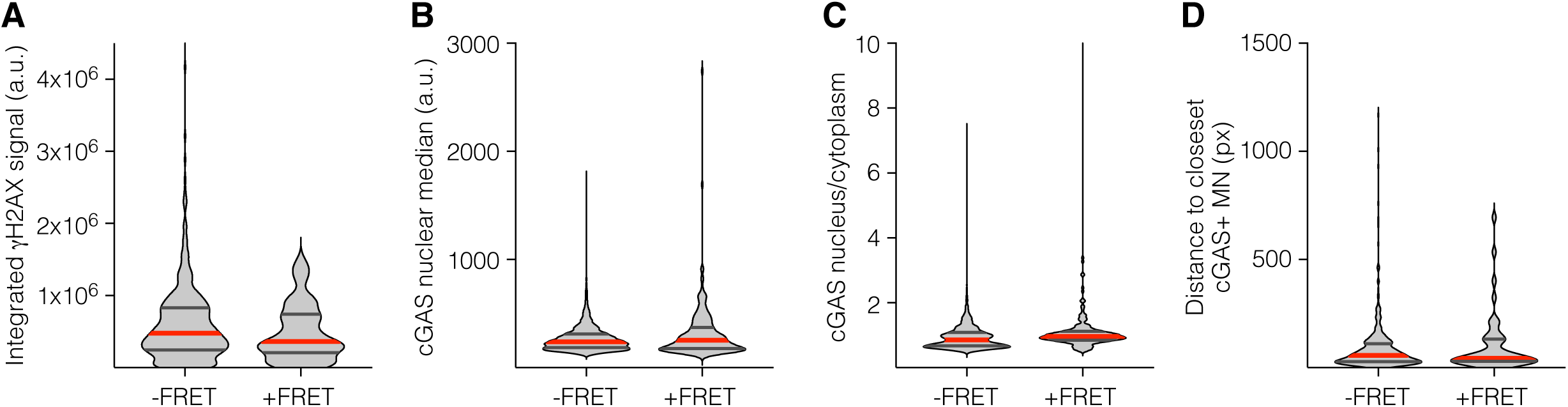
Analysis of DNA damage and cGAS localisation parameters following IR. (A-D) Correlative analysis of cGAMP reporter FRET and immunofluorescence. Cells were imaged for FRET under live conditions 48 h after treatment with 15 Gy of IR. Subsequently, cells were fixed, stained with the indicated antibodies, and corresponding cells were identified in each dataset. Violin plots show the general distribution, with the median indicated by a red line, and the interquartile range indicated by black lines. (A) Analysis of gH2AX levels in cells with and without FRET. (B) Analysis of median cGAS intensity in cells with and without FRET. (C) Analysis of nuclear/cytoplasmic ratio of cGAS in cells with and without FRET. (D) Distance to closest cGAS-enriched micronucleus in cells with and without FRET.

**Figure S5.**
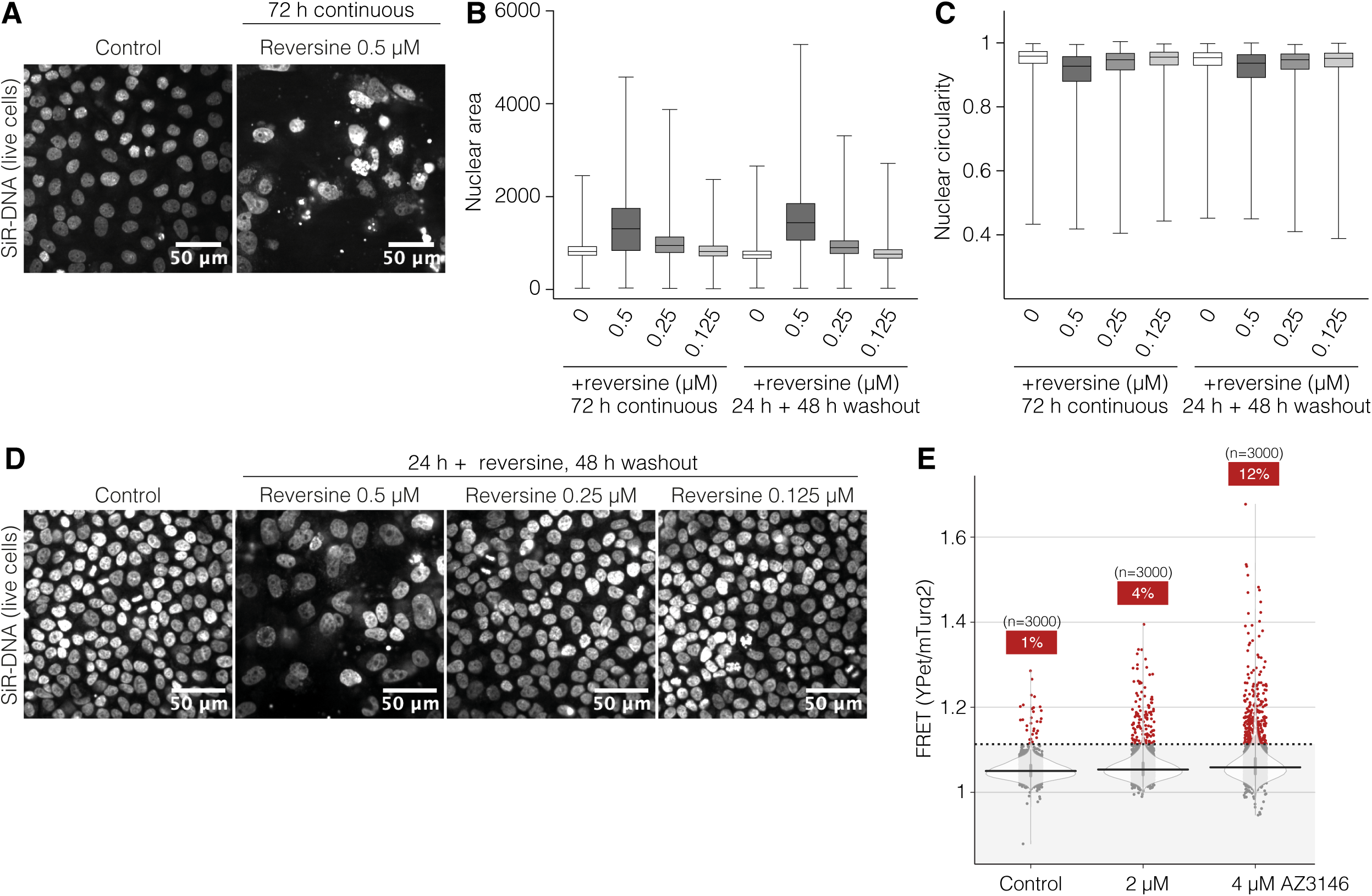
Further characterisation of the relationship between micronuclear cGAS and its activation. (A-D) Nuclear abnormalities induced by MPS1i in MCF10A cells. Cells were treated with reversine at the indicated dose for either 72 h, or for 24 h followed by washout and 48 h chase. (A) Representative images of MCF10A nuclei treated with 0.5 µM reversine or DMSO control for 72 h. SiR-DNA staining imaged from live cells. (B-C) Quantification of nuclear shape (as determined from Hoechst immunofluorescence staining) following reversine treatment, using either nuclear area (B) or circularity (C) as a marker. Box and whisker plots are shown with, centre lines indicating the median, box boundaries indicating upper and lower quartiles, and whiskers indicating maximum and minimum. (D) Representative images of MCF10A nuclei treated with the indicated dose of reversine or DMSO control for 24 h, followed by washout and 48 h chase. SiR-DNA staining imaged from live cells. (E) MCF10A stably expressing the cGAMP reporter were treated for 72 h with the indicated dose of AZ3146, and subsequently imaged for FRET ratio measurement. Dots represent values from individual cells. Violin plots show the general distribution, with the median indicated by a horizontal line, and the interquartile range indicated by a grey vertical line. Red dots indicate cells containing cGAMP (as defined by a FRET response higher than 99% of control cells). Numbers in red boxes indicate the percentage of cells containing cGAMP.

**Movie S1.**

Time-lapse microscopy of MCF10A cells stably expressing the cGAMP reporter treated with DNA transfection, DMXAA (10 µg/mL) or diABZI (1µM). Related to Figure 2A-B.

**Movie S2.**

Time-lapse microscopy of MCF10A cells stably expressing the cGAMP reporter activating cGAS after 7Gy irradiation. Related to Fig 3F.

**Movie S3.**

Time-lapse microscopy of mScarlet-cGAS being recruited to MN in MCF10A cells stably expressing both the cGAMP reporter and mScarlet-cGAS treated with reversine 0.5 µM. Related to Fig 5F-G

## Methods

### Cells

MCF10A cells were cultured in DMEM/F12 supplemented with 5% (v/v) horse serum (Thermo Fisher Scientific), 20 ng/ml EGF (Sigma-Aldrich), 0.5 µg/ml hydrocortisone (Sigma-Aldrich), 100 ng/mL cholera toxin (Sigma-Aldrich), 10 µg/mL insulin (Sigma-Aldrich) and 1% penicillin-streptomycin (Gibco). SUM149PT cells were cultured in Ham’s F12 medium (Gibco) supplemented with 5% (v/v) heat inactivated foetal bovine serum (FBS) (Gibco), 10 μg/ml insulin (Sigma), 0.5μg/ml hydrocortisone (Sigma) and penicillin/ streptomycin (Gibco). HeLa and HEK293FT cells were grown in DMEM supplemented with 10% FBS and 1% penicillin-streptomycin (Gibco).

MCF10A stably expressing the cGAMP reporter, the cGAMP-blind reporter, GFP-IRF3 or mScarlet3-cGAS were generated by lentivirus transduction. To generate lentiviral particles, pLex306 plasmid (David Root, Addgene plasmid #41391) containing the open reading frames of either sequence was co-transfected with psPAX2 and pMD2.G in HEK293FT cells at 50% confluency in a T75 flask using lipofectamine 2000 (Invitrogen). Media was changed after 24 h, and supernatant was collected after 3 days and centrifugated to remove cell debris. Supernatant was mixed 3:1 with 4x concentrator solution (80 g PEG-8000, 1 4g NaCl in 200 ml 1x PBS, pH adjusted to 7.0∼7.2) and mixed overnight by constant rotation at 4 °C. The mix was centrifuged at 1600 g (60 min, 4 °C), the pellet was resuspended in PBS and used to transfect MCF10A in T25 flasks. Successfully transduced clones were selected by flow cytometry based on expression level.

SUM149PT cells stably expressing the cGAMP reporter and the cGAMP-blind reporter were generated by lentivirus transduction. To generate lentiviral particles, pLex306 plasmid (David Root, Addgene plasmid #41391) containing the open reading frames of either was co-transfected with psPAX2 and pMD2.G in HEK293FT cells at 50% confluency in a 10 cm dish using lipofectamine 2000 (Invitrogen). Media was changed after 24 h, and the supernatant was collected after 3 days, filtered with a 0.4 μm pore filter to remove cell debris, and was used to infect cells in suspension, in a 3cm dish. The infected cells were then expanded, sorted by flow cytometry and pools of transduced cells were generated.

Hela H2B-RFP cGAS knock-out and EGFP-cGAS rescue cells were used as previously described^8^.

### Reagents

Chemical compounds dissolved in sterile DMSO were used at the following concentration unless indicated otherwise: DMXAA STING agonist (Invivogen, 10 µg/mL), cGAS inhibitor (G150, Selleckchem, 10 µM), Reversine (Fisher Scientific, 0.5 µM), AZ3146 (Fisher Scientific), Doxorubicin (Medchem Express). Chemical compounds dissolved in sterile water were used at the following concentration unless indicated otherwise: 2’3’-cGAMP (Invivogen, 10 µg per electroporation with 100,000 cells), diABZI compound 3 (Invivogen, 1µM).

### Antibodies

cGAS was detected with #15102 from Cell Signaling Technology (CST) (1:300 dilution for immunofluorescence; 1:1000 dilution for Western Blot), total IRF3 with #4947 from CST (1:500 dilution for IF), phospho-IRF3 Ser396 with #4947 from CST (1:100 for IF), phospho-STAT1 Tyr701 with #9197 from CST (1:300 dilution for IF), yH2AX Ser139 with 05-636-25UG from Millipore (1:500 for IF), STING with 13647S from Cell Signalling Technology (1:1000 for Western Blot), α-tubulin with T9026 from Sigma (1:5000 for Western Blot).

### Western blotting

Cell pellets were harvested and lysed in 1x RIPA buffer (Merck 20-188) supplemented with 1x protease inhibitor cocktail (Roche 11873580001). Samples were diluted in 4x LDS sample buffer (Life Technologies NP0008) containing 1x sample reducing agent (Life Technologies NP0009) and boiled for 5min at 95 °C prior to being subjected to gel electrophoresis and western blotting. The Odyssey Infrared Imaging System (Li-Cor M) was used for detection.

### Plasmid construction

To generate a FRET-based cGAMP reporter, PCR amplification of the coding sequences of mTurquoise2^36^, mSTING fragment^35^ and YPet^37^ were cloned by restriction digest on an ePB piggyBac backbone^70^. For the cGAMP-blind reporter, the mSTING fragment was replaced with the Y239S/T262A mutant^35^. To generate lentiviral plasmids containing the cGAMP reporter, a fragment containing mTurquoise2 and the mSTING CDN was amplified by PCR and cloned in a pLex306 backbone (David Root, Addgene plasmid #41391). The sequence of E. coli codon-optimised YPet from^44^ was synthesised as a gBlock (IDT) and cloned onto the C-terminus of mSTING.

The GFP-IRF3 construct was made by PCR-amplifying GFP-IRF3 from^8^ and cloning it into the lentiviral pLex306 backbone. The mScarlet3-cGAS (hereafter mScarlet-cGAS) construct was made by PCR-amplifying plasmid containing human cGAS for protein expression^8^. The mScarlet3 was synthesized by IDT. These DNA fragments were inserted into the lentiviral pLex306 backbone.

### Reporter protein expression and purification

Wild type and mutant reporters were expressed in Rosetta (DE3) pLysS cells from pET15b plasmid backbones with an N-terminal 6xHis tag. Cells were seeded into terrific broth (24 g/l yeast extract, 12 g/l bacto tryptone, 0.004% glycerol, 17 mM KH_2_PO_4_, 72 mM K2HPO4) containing 0.4% glucose, 50 µg/ml carbenicillin and 34 µg/ml chloramphenicol, and incubated shaking at 37 °C until an OD_600_ of ∼0.8. Expression was induced with 0.25 mM IPTG at 18°C for 6 h. Cleared lysate was incubated with HisPur Ni-NTA resin (Fisher Scientific 10038124) for 1 hr at 4 °C, and recombinant protein was eluted using elution buffer (50 mM Tris-Cl pH 7.5 @ 4 °C, 200 mM NaCl, 300 mM imidazole, 5 mM β-Mercaptoethanol (BME). Eluted fractions were pooled and further purified on a 1 ml HiTrap FF Q column (VWR 17-5053-01) using an AKTA FPLC system (Cytiva). Flowthrough fractions were collected and concentrated before size exclusion chromatography on a Superdex 200 10/300 GL gel filtration column (Cytiva) using gel filtration buffer (40 mM Tris-Cl pH 7.5, 100 mM NaCl, 0.5 mM TCEP). The purified reporters were diluted in gel filtration buffer containing glycerol to a final concentration of ∼40% glycerol and stored at -20 °C.

### Mass Photometry

All measurements were perfomed on an OneMP mass photometer (Refeyn Ltd, Oxford, UK). Glass slides (High Precision microscope cover glasses 24 x 50 mm, Marienfeld 630-2186) were cleaned with water and isopropanol four times and dried using compressed air. A drop of emersion oil (Olympus Immoil-F30CC) was placed on the objective and a glass slide with a gasket (Reuseable Culturewell gasket 3 mm diameter x 1 mm depth, Grace Bio-Labs 103250) adhered to the glass was placed on top of the emersion oil. The instrument was calibrated by addition of 15 µl of protein standards from Native Mark Unstained Protein Standard (Thermo Fisher LC0725) in buffer A (10 mM Tris-HCl pH 7.4, 10 mM KCl, 1.5 mM MgCl_2_). For calibration, the objective was focused through the buffer-free method. For cGAMP reporter readings, 100 nM reporter pre-incubated with 250 nM cGAMP for 15 min at 25 °C or 100 nM reporter control reaction was prepared. The objective was focused using the drop-wise dilution method by addition of 13 µl of buffer A and 2 µl of corresponding reactions. Upon gentle mixing, immediately the mass photometry recording was started in Acquire MP. Recordings were done in the regular field of view for 60 seconds, with 1000 to 4000 particle landing events per movie. Analysis of each movie was performed with the mass photometry software, Discover MP. Masses of the major forms of reporter +/- cGAMP were estimated by fitting a Gaussian distribution into mass histograms and taking value at the mode of distribution.

### Cell lysate preparation for cGAMP measurement

cGAS knockout and rescue HeLa H2B-RFP cells ^8^ were plated at 600,000 cells/well in a 6 well tissue culture dish in DMEM plus 100 ng/ml of doxycycline. 24 h later, media was replaced with fresh media without antibiotics and cells were transfected with a cocktail of 2.6 µg pBlueScript II Sk (+), 8.62 µl ViaFect (Promega E4981) and 103 µl of Opti-MEM (Gibco 31985062). 8 hr later, 1.2 x 10^6^ cells were harvested following trypsinisation and two rounds of washes with 1x PBS. Samples were processed using a previously described procedure^71^. Briefly, each cell pellet was resuspended in 200 µl 80:20 methanol:water and incubated overnight at -80 °C. Subsequently, samples were subjected to 2 rounds of vortex, freeze/thaw cycles in liquid nitrogen, sonicated in an ice water bath for 10 min and clarified by centrifugation at 21,000 x g for 20 min at 4 °C. Extracts were dried using a speed vac and reconstituted in 50 µl of buffer A (10 mM Tris-HCl pH 7.4, 10 mM KCl, 1.5 mM MgCl_2_).

### cGAMP measurement in cell lysates

500 nM reporter concentrations were used to measure cGAMP produced in cell lysates. Reconstituted cell lysate was incubated with the reporter for 15 min at 25 °C and transferred into 96-well half area black flat bottom plates (Sigma-Aldrich CLS3694-25EA). FRET was measured on a POLARstar Omega (BMG Labtech) with mTurqoise2 detected with 430 nm (excitation)-480 nm (emission) filter pairs, and YPet-FRET detected with 430 nm (excitation)-530 nm (emission) filter pairs. Gain adjustments were done for reporter without cGAMP for 430-480 nm and for reporter with saturating concentrations of cGAMP for 430-530 nm. A standard curve was generated for each experiment.

### Native gel electrophoresis

1 µg of purified reporter +/- 2.64 µM cGAMP were incubated for 15 min at 25 °C. Samples were mixed with 2 M sucrose at a 5:1 volume ratio, and resolved on Mini-Protean TGX 4-15% gels (Bio-Rad 4561086) in a buffer containing 25mM Tris and 192 mM glycine (pH 8.3) at room temperature (1h, 100 V). The gel was stained with GelCode Blue Safe Protein Stain (Thermo Fisher Scientific 10763505), and imaged on an LI-COR Odyssey M infrared scanner.

### Analysis of CCL5 expression

For irradiated samples, ∼2×10^5^ cells were seeded in a well of a 6-well plate. Next day, cells were irradiated with 15Gy. Immediately after irradiation, media was replaced with fresh media. Cells were harvested for RNA extraction 48 h post irradiation. For DNA transfection samples, 4×10^5^ cells were seeded in a well of a 6-well plate. Next day, media was removed from cells and replaced with 0.7 ml of media containing no antibiotics. To generate the transfection mix, 2 μg of Herring-testis DNA (Sigma-Aldrich, # D6898-1G) were combined with 0.15 ml Opti-MEM (Life Technologies # 51985-034). Separately, 6 μl Lipofectamine 2000 (Life Technologies, # 11668) were mixed with 0.15 ml Opti-MEM. Both mixtures were incubated at room temperature for 5 min, combined and incubated at room temperature for 15 min before being added to the cell dish. Samples were harvested for RNA extraction 6 h post stimulation.

To prepare cDNA, ∼ 4×10^5^ cells were harvested and resuspended in 0.8 ml TRI-Reagent (Sigma-Aldrich, # T9424-25ML). This mixture was frozen at −80 °C until use. For RNA extraction, 0.2 ml chloroform were added after thawing, and samples were transferred to Phase-lock gel tubes, heavy (VWR, # 733-2478). Following extensive mixing, samples were incubated at room temperature for 2 min before centrifugation for 5 min (12.000 g; 4 °C). The aqueous phase was removed and mixed with an equal volume of ethanol. After mixing, samples were loaded onto RNeasy columns (Qiagen), washed once with buffer RW1 and twice with buffer RPE. Samples were eluted using 35 μl of nuclease-free water. ∼1 μg RNA of each sample was used to generate cDNA using the High-Capacity cDNA Reverse Transcription Kit (Applied Biosystems, # 4368813) according to the manufacturer’s instructions. cDNA samples were diluted 1:10 in nuclease-free water, and 4 μl of each cDNA reaction were used for quantitative PCR with the SYBR™ Green PCR Master Mix (Applied Biosystems, # 4344463), carried out on an StepOnePlus Real-Time PCR System (Applied Biosystems, # 4376600). Amplification conditions were 95 °C 10 min → 40x (95 °C 15 sec ® 60 °C 1 min). Oligos were AATCCCATCACCATCTTCCA and TGGACTCCACGACGTACTCA for GAPDH^72^, and TGCCCACATCAAGGAGTATTT and CTTTCGGGTGACAAAGACG for CCL5^73^.

### FRET flow cytometry

HEK293FT cells were seeded in 10 cm dishes to be at 80% confluency on day 1. The cells were transfected with the cGAMP reporter using Lipofectamine 2000 (Invitrogen) as per the manufacturer’s instructions and were treated 24 h later with either media or diABZI (1 μM) for 1.5 h. Cells were then washed in PBS, trypsinised, spun down and the pellet resuspended in PBS.

Live cell flow cytometry FRET measurements were performed on a BD FACSymphony S6 cell sorter (BD Biosciences). mTurq2 was excited using a Violet 405 nm diode laser with a power of 100 mW. mTurq2 was detected with the 405-474/25 channel at 510 V. YPet-FRET was detected with the 405-530/30 channel at 400 V. Data was analysed using FlowJo. Briefly, after gating for single cells based on FSC-H and FSC-A, we gated the mTurq2+ population based on the parental cells that did not express the reporter.

### cGAS stimulation with DNA in cells

For experiments using DNA in cells as a stimulus for cGAS activation, cells were seeded on a µCLEAR 96 well plate (Greiner Bio-one, 655090) in MCF10A culture media without phenol red to be at 80% confluency on the day of imaging. For each well, pBluescript SK(+) plasmid DNA was suspended in 25 µL MCF10A culture media without serum or antibiotics, mixed with a second tube containing 0.5 µL Lipofectamine 2000 (Invitrogen) in 25 µl MCF10A culture media without serum or antibiotics, left to incubate for 20 min, and then added to the well containing the cells in 100 μl of MCF10A culture media. Unless specified otherwise, cells were imaged 8h post-transfection.

### Live cell imaging

For live cell experiments, MCF10A cells stably expressing the cGAMP FRET reporter were seeded on a µCLEAR 96 well plate (Greiner Bio-one, 655090) or µ-Slide 8 Well High ibiTreat #1.5 polymer tissue-culture treated chamber slide (IBIDI) in MCF10A culture media without phenol red to be at ∼80% confluency on the day of imaging. For radiation experiments and unless specified otherwise, cells were exposed to 15Gy of X-ray radiation and let to recover for 48 h before imaging. Nuclei were stained with 200 nM SiR-DNA (Spirochrome) 2-6 h before imaging.

For FRET experiments in HeLa cells, 100,000 cGAS knockout HeLa cells were transiently co-transfected with the cGAMP reporter (or cGAMP-blind mutant) with or without 10 µg cGAMP using the Neon transfection system (Invitrogen, 10 µl tips, program: 1005 V, 2 pulses, 35 ms). Following transfection, cells were immediately seeded on µ-Slide 8 Well High ibiTreat #1.5 polymer tissue-culture treated chamber slide (IBIDI) containing 300 μl DMEM without phenol red + 5% FBS. Cells were let to recover for 48 h, then imaged.

FRET images were acquired on a Zeiss Axio Observer Z1 equipped with a Yokogawa CSU-X1 spinning disk unit, as well as with 445 nm 75mW, 488 nm 50 mW, 514 nm 40 mW, 561 nm 50 mW and 640 nm laser illumination sources. Emission filters (Semrock) were as follow: 440/521/607/700, ChromaZET440/514/561/640, 482(35), 542(27), 617(73), 525(30). To maximise the number of cells in each field of view, a Zeiss Plan-Apochromat 20x/0.8 objective was used for all experiments. Temperature and CO_2_ were maintained at 37 °C and 5% CO_2_ by an Okolabs Bold Line T-Unit1 and Okolabs Bold line CO_2_/O_2_ unit.

To assess bleedthrough between the CFP and YFP FRET channels, cGAS knockout HeLa cells were transfected with an mTurquoise2-mSTING-mTurquoise2 construct, or a H2B-Citrine construct. Using the CFP excitation laser, 39.7±3.1% (mean ± SD) of the mTurquoise2-mSTING-mTurquoise2 fluorescence could be detected with the YFP emission filter, and 0% of the H2B-citrine signal could be detected with the CFP emission filter. When using the YFP excitation laser, no mTurquoise2-mSTING-mTurquoise2 fluorescence could be detected with either the CFP or YFP emission filters, and no H2B-citrine signal could be detected with the CFP emission filter.

Snapshots images were acquired on a Hamamatsu Flash4.0 V3 with full laser power and 2×2 camera binning. Exposure was set at the same 600ms value for all three FRET channels (CFP, YFP and CFP-excited YFP emission) and at 100 ms for SiR-DNA staining. To capture rare FRET activation events following DNA damage induction, at least 50 to 80 fields of view were acquired per well and multiple technical replicates were combined per experimental condition.

For time lapse FRET, mScarlet3-cGAS and GFP-IRF3 imaging, the camera was switched to a Photometrics Prime95b with Gain 3 and laser power reduced to 20% to limit photodamage. Images were acquired every 15 or 20 minutes for FRET experiments, and every 5 minutes for GFP-IRF3. For the cumulative IR FRET activation plot in Figure 4G, images were acquired every 120 min on a Flash4 camera with full laser power. For timelapse FRET DNA damage experiments, the acquisition was started either immediately following X-ray irradiation, or 3 h hours after 0.5 µM Reversine addition.

When drugs were added during a timelapse acquisition, they were diluted to a 10X concentration in 10 µl imaging media, then added dropwise to the well containing cells in 100 µl imaging media. Care was taken not to touch the plate with the pipette tip and not to open the imaging chamber for more than a minute to limit temperature and CO_2_ shifts. For stimulation with DNA during a timelapse, 50 µl of transfection mix (see section ‘cGAS stimulation with DNA’) in MCF10A imaging media without serum or antibiotics were added dropwise to the well, putting the total well volume to 150 µl.

### Fixed cell FRET imaging

Cells seeded in µCLEAR 96 well plate (Greiner Bio-one 655090) were treated with 200 nM Sir-DNA for 2-4 h, washed in PBS once then fixed 10 min in 4% paraformaldehyde (PFA). Cells were washed 3x in PBS then stored protected from light at 4 °C. Cells were imaged as described above for snapshots of live cells.

### Immunofluorescence

Cells seeded in µCLEAR 96 well plate (Greiner Bio-one 655090) were washed with PBS then fixed in 4% PFA for 10 min, washed 3 times in PBS then stored at 4 °C for further processing. Cells were permeabilised in PBS+0.1% triton for 10 min, washed in PBS and blocked 1 h in 3% BSA. Cells were incubated 1 h in primary antibody, washed 3x in PBS+0.1% tween (PBST) and incubated for a further 1 h with secondary antibody. Cells were then washed 3x in PBST, incubated 5 min in 10 µg/ml Hoechst 33342 (Thermo) then stored in PBST at 4 °C.

Immunofluorescence images were acquired on an ImageXpress (Molecular devices) equipped with (excitation filters:) 377/54, 475/34, 531/40, and 631/28, as well as (emission filters:) 447/60, 536/40, 593/40, 624/40, 692/40. Images were acquired with a 20x objective in widefield mode.

### Image processing and data analysis

Segmentation, Tracking and pixel value quantification were performed using custom Jython scripts run in FiJi^74^. Further processing was performed with python scripts based on Pandas and Numpy for data processing and matplotlib and seaborn to generate graphs. Alternatively, processed data was exported as a table to be imported and plotted in Graphpad Prism. All code used in this work is available on Github.

For both live and fixed cells, nuclear signal (SiR-DNA, H2B-RFP or Hoechst 33342) was used to segment nuclei in FiJi using StarDist^75^. A 3 px wide ring was generated 3 px away and around the periphery of every nuclei to obtain a ‘cytosolic’ region. FRET ratios were calculated by dividing measured pixel values in the YPet FRET channel (CFP excitation laser, YFP emission filter) by the measured pixel values in the mTurquoise2 Donor channel (CFP excitation laser, CFP emission filter). Care was put into determining which approach gave the most robust to noise and reproducible measurements across a range of conditions, and it was ultimately decided to measure median values of channels of interest in the cytosolic region. All FRET data in this work is showing cytoplasmic FRET ratio values.

Despite being derived from a clone, slight variability in expression levels are apparent in cells stably expressing the cGAMP FRET reporter and FRET measurements do not appear to be fully accurate below a certain expression level. To account for that, the YFP fluorescence channel (YFP excitation laser, YFP emission filter) was used to assess expression level and cells in which reporter expression was deemed too low were fully excluded from any further analysis. In this work, a threshold of 1500 (au) for median YFP cytosolic intensity was empirically determined to be the most robust and reproducible across different treatment conditions (Figure S3A). This threshold needs to be optimised depending on cell line, expression level and imaging conditions.

For all FRET experiments showing heterogeneous and partial response to DNA damage, a threshold of activation was defined in each experiment as the 99th percentile of all measured FRET values in the control condition. Any cell for which the measured FRET ratio was above this threshold was classified as active and plotted as a red dot, while cells below this threshold were plotted as a grey dot. The percentage of cells in the population which is above the threshold (and therefore classified as ‘active’) is indicated for each condition at the top of the graph in a red box, along with total cell number (active or inactive) in the given condition.

Treatments such as radiation and high doses of reversine severely impact cell growth compared to untreated control condition. To display comparable cell numbers across all conditions and when applicable, cells were randomly selected using the unbiased “sample” function from Python pandas before plotting. Percentage of active cells are calculated from the total population (before sampling).

For figure S3B, significant autofluorescence from doxorubicin affected the baseline of measured FRET ratio. To account for that non-cGAMP related shift, the measured FRET ratio of each cell was divided by the average FRET measurement of MCF10A cells treated with the same doxorubicine concentration but stably expressing the mutant cGAMP-blind reporter instead of the WT one. Figure S3B is the only graph in this work that was normalised to average cGAMP-blind reporter FRET values.

For time lapse experiments, detected nuclei from each frame were linked in time using TrackMate^76^. To account for cells entering or exiting the field due to cell motility, only traces that lasted more than 37.5h were considered for analysis in Fig 3E. For timelapse experiments, individual traces were smoothed with a rolling median algorithm using a centred 1h window.

To automatically detect cGAS+ micronuclei in immunofluorescence, the cGAS immunofluorescence staining channel was segmented using StarDist. Detected particles were filtered based on size and fluorescence to exclude main nuclei and only keep cGAS+ MN. Distances from every detected cGAS+ particle to all neighbouring main nuclei (defined by the Hoechst staining) were computed, and a Gale-Shapley algorithm used to match every detected cGAS+ particle to its closest main nucleus.

### Correlation of live FRET imaging and immunofluorescence

Following acquisition of live cell FRET images, cells were immediately washed in PBS then fixed in 4% PFA for 10 minutes. Immunofluorescence incubations were performed as described above. To limit interaction with CFP and YFP channel, only a single primary antibody was incubated with cells, and further detected by a red fluorescence Cy3 secondary antibody.

For all wells in which FRET images were previously acquired, a tiled mosaic encompassing the full well was generated and the Hoechst 33342 and Cy3 immunofluorescence staining acquired on either a widefield ImageXpress (Molecular devices) with a 20x objective, or on a spinning disk confocal Opera Phenix (Perkin-Elmer) with a 40x water-immersion objective.

To identify cells from the live imaging experiment in the follow-up immunofluorescence acquisition, individual images from the live FRET experiment were resized to account for differences of pixel size and magnification between the two imaging setups. Live cell images and the full well immunofluorescence mosaic montage were segmented using StarDist to generate binary images, which were further processed to obtain their Fourier transform. For each field from the live imaging, the associated Fourier transform conjugate was multiplied with the mosaic image Fourier transform, then converted back with an inverse Fourier transform to compute the location of the FRET live imaging field in the follow up immunofluorescence full well mosaic. Segmented objects in both live imaging and IF datasets were filtered by size to exclude debris, and XY coordinates of every cell from either the live imaging or immunofluorescence images were merged into a common coordinate system adjusted for magnification and pixel-size differences. Distances between every cell from the FRET live cell experiment to every cell from the immunofluorescence acquisition were computed into a cost matrix which was resolved using SciPy linear sum assignment algorithm to identify every cell from the live imaging experiment in their immunofluorescence counterpart. Small images centred on the nuclei (SiR-DNA vs Hoechst) of identified corresponding cells were generated to assess matching quality.

Image analysis of the immunofluorescence data was performed on each individual tiles instead of the reconstituted mosaic montage to prevent potential stitching artefacts from affecting the pixel values. Cells from the individual IF tiles were matched to the mosaic montage and the corresponding live cells as described in the previous paragraph.

## Author contributions

CZ and VL designed the study. VL designed the cGAMP reporter and carried out and analysed the majority of experiments. NA performed and analysed biochemistry; LD and VL carried out the flow cytometry analysis of FRET; FT carried out the QPCR analysis induction; TK and VL carried out the time-lapse analysis of cGAS micronuclear enrichment and cGAMP production. AK generated and analysed the SUM149PT cells expressing the reporter. CZ and VL wrote the manuscript, with contributions from all the authors.

## Acknowledgments

Work in the laboratory of CZ is supported by Cancer Research UK (RCCFEL\100092), by the Cancer Research UK Radiation Research Centre of Excellence at The Institute of Cancer Research and The Royal Marsden NHS Foundation Trust (A28724), and by Breast Cancer Now (2023.05PR1625). TK is supported by AMED (JP23ad0027013). We thank Joshua Woodward for providing the original cGAMP FRET reporter; Chris Lord for SUM149PT cells, Kai Betteridge, Ana Stojiljkovic, Queenie Lai, Ross Scrimgeour from the ICR light microscopy facility for help with microscopy; Stephen Hearnshaw for help with mass photometry; Hira Ale, Matt Guelbert and Yuliya Semochkina from the ICR flow-cytometry facility for help with flow-cytometry. We also thank the members of the Zierhut lab and Max Douglas for critical reading of the manuscript, and members of the Zierhut lab, the Mansfeld and Douglas labs at ICR for helpful discussion.

